# Targeting NGF but not VEGF or BDNF signaling reduces endometriosis-associated pain in mice

**DOI:** 10.1101/2023.12.05.570181

**Authors:** Tiago H. Zaninelli, Victor Fattori, Olivia K. Heintz, Kristeena R. Wright, Philip R. Bennallack, Danielle Sim, Hussain Bukhari, Kathryn L. Terry, Allison F. Vitonis, Stacey A. Missmer, Avacir C. Andrello, Raymond M. Anchan, Stephen K. Godin, Dara Bree, Waldiceu A. Verri, Michael S. Rogers

**Author notes:** Corresponding author Address: Vascular Biology Program, Boston Children’s Hospital, Department of Surgery, Harvard Medical School, 11.211 Karp Family Research Bldg, 300 Longwood Ave, Boston, MA 02115, United States. E-mail address (M.S. Rogers).

## Abstract

**Introduction:** Endometriosis is a chronic inflammatory disease that affects ∼10% of women. A significant fraction of patients experience limited or no efficacy with current therapies. Tissue adjacent to endometriosis lesions often exhibits increased neurite and vascular density, suggesting that disease pathology involves neurotrophic activity and angiogenesis.

**Objectives:** We aim to evaluate the potential for key tyrosine-kinase-receptor-coupled neurotrophic molecules to contribute to endometriosis-associated pain in mice.

**Methods:** The levels of VEGFR1 regulators (VEGFA, VEGFB, PLGF, and sVEGFR1) were quantified by ELISA in peritoneal fluid from endometriosis patients undergoing surgery and used to calculate VEGFR1 occupancy. We used genetic depletion, neutralizing antibody, and pharmacological approaches to specifically block ligand (NGF or BDNF) or neurotrophic receptor (VEGFR1, TRKs) in a murine model of endometriosis-associated pain. Endometriosis-associated pain was determined using the von Frey filaments method, quantification of spontaneous abdominal pain-related behavior, and thermal discomfort. Diseases parameters were evaluated by lesion size and prevalence.

**Results:** We found that entrectinib (pan-Trk inhibitor) or anti-NGF treatments reduced evoked pain, spontaneous pain, and thermal discomfort. In contrast, even though receptor occupancy revealing that VEGFR1 agonist levels are sufficient to support pain, blocking VEGFR1 signaling via antibody or tamoxifen-induced knockout did not reduce pain or lesion size in mice. Targeting BDNF-TrkB with an anti-BDNF antibody also proved ineffective.

**Conclusions:** This suggests NGF-TrkA signaling, but not BDNF-TrkB or VEGF-VEGFR1, mediates endometriosis-associated pain. Moreover, entrectinib blocks endometriosis-associated pain and reduces lesion sizes. Our results also indicated that entrectinib-like molecules are promising candidates for endometriosis treatment.

**Credit author statement:** **Conceptualization:** T.H. Zaninelli, V. Fattori, and M.S. Rogers; **investigation and data curation:** T.H. Zaninelli, V. Fattori, O.K. Heintz, K.R. Wright; A.C. Andrello, W.A. Verri Jr, M.S. Rogers; **funding acquisition:** M.S. Rogers,, S.A. Missmer, R.M. Anchan; **methodology:** T.H. Zaninelli, V. Fattori, and M.S. Rogers; **human sample collection:** S.A. Missmer, A.F. Vitonis, K.L. Terry, R.M. Anchan; **animal breading and VEGFR1 ablation:** D. Sim and H. Bukhari; **resources:** A.C. Andrello; D. Bree, T. Zheng, J. Wagner, W.A. Verri Jr, and M.S. Rogers; **project administration:** T.H. Zaninelli; **supervision:** V. Fattori, W.A. Verri Jr, and M.S. Rogers; **visualization:** T.H. Zaninelli, V. Fattori, W.A. Verri Jr, and M.S. Rogers; **writing–original draft:** T.H. Zaninelli; **writing – editing and reviewing:** all authors. All authors have read and approved the final version of the manuscript.

## Introduction

Endometriosis is an estrogen-dependent inflammatory disease characterized by the presence of endometrium-like tissue in the abdominal cavity or pelvic space^1^. The disease affects 10% of people born with a uterus^2^. The most studied pathway for endometriosis genesis is retrograde menstruation^3–5^. During this process, danger-associated molecular patterns (DAMPs)^6^ activate resident immune cells and trigger inflammation^7^. The inflammatory process is orchestrated by pro-inflammatory mediators and growth factors secreted by activated immune cells^7,8^. The growth-factor-enriched inflammatory milieu favors angiogenesis and neurogenesis, resulting in vascularized and innervated endometriotic lesions teeming with activated immune cells^9^. This inflammation is key to pain, which is a prevalent clinical feature of those diagnosed with endometriosis^3,4^.

Endometriosis-associated pain appears as chronic pelvic pain, dysmenorrhea, dyspareunia, and dyschezia, affecting the social and professional quality of life and mental health of patients with endometriosis^10^. For instance, patients with endometriosis lose approximately 10 hours of work weekly, because of reduced effectiveness during work due to endometriosis’symptons^11^. Currently, endometriosis-associated pain treatment relies pharmaceutical and nonpharmaceutical approaches including the use of non-steroidal anti-inflammatory drugs (NSAIDS), opiates and other analgesics, hormonal therapy, pelvic therapy, acupuncture, and the surgical removal of lesions^12^. Nevertheless, current treatments show limited efficacy in reducing pain and often present unwanted side effects^13^. Importantly, up to 30% of patients with endometriosis do not respond to the current therapies^14^. Therefore, there is an unmet need to develop or repurpose effective and safe drugs for the treatment of endometriosis-associated pain.

The endometriotic microenvironment is associated with increases in multiple tissue remodeling growth factors, including nerve growth factor (NGF) and vascular endothelial growth factor (VEGF), which are important mediators of neurogenesis and angiogenesis, respectively. In addition to their growth-related role, those mediators are also described to be involved in pain^15,16^. Strong evidence shows that lesion growth is dependent on VEGFR2-mediated angiogenesis^17–20^, while VEGFR1 contributes to inflammatory milieu maintenance^21^. Moreover, VEGF has shown promise in a panel of biomarker candidates for non-invasive endometriosis diagnosis^22^. In cancer models, tumor-derived VEGF-A/B, and PlGF-2 increase pain sensitivity by activating VEGFR1^16^. Similarly compelling evidence demonstrates that NGF might be involved in endometriosis pain^23^. High expression of *NGF* in deep adenomyotic nodules is correlated to hyperalgesia^13^ and both *NGF* and *NTRK1* (TRKA) expression in human lesions correlate with deep dyspareunia in women with endometriosis^24^. Furthermore, a recent GWAS study showed that a variant in *NGF* is associated with migraine and dysmenorrhea for women with endometriosis^25^. However, the relative contributions of various neurotrophic factors to endometriosis-associated pain remain unclear.

To evaluate the effect of modulating cellular signaling of the neurotrophic molecules VEGF, NGF, and BDNF as potential therapeutic targets for endometriosis-associated pain, we used our validated mouse model of endometriosis. This model mimics neuronal and behavioral changes consistent with the disease phenotype in women^26^. Moreover, resulting lesions exhibit features that resemble human lesions such as the presence of nerve fibers, glands, and immune cells. The model responds to clinically active drugs, and therefore, might be useful for finding novel or repurposed therapies^26^.

## Materials and Methods

### Patient samples

Samples (n=33) of peritoneal fluid (PF) were collected and processed as part of the Women’s Health Study: From Adolescence to Adulthood (A2A) cohort^27^. Samples were collected following the WERF EPHect protocol^28^ with some deviations. PF was housed in an incubator immediately after collection until the specimen could be transported to the lab for processing. Samples were centrifuged at 300g for 10 minutes at 4°C and frozen at -80°C using a Mr. Frosty freezing container. Samples were kept frozen until aliquoting and only freeze-thawed twice. Additional samples (n=9) were similarly collected following the WERF EPHect protocol from patients undergoing exploratory laparoscopy surgery for endometriosis at Brigham and Women’s Hospital (Boston, MA, USA).

### Animals

Healthy and immunologically competent C57BL/6 (8 weeks old, 20-25g, female, strain # 664 [RRID:IMSR_JAX:000664]), B6.Cg-Flt1^tm1.1Fong/^J (8 weeks old, 20-25g, male and female, Vegfr-1^flox^, strain # 28098 [RRID:IMSR_JAX:028098]), and B6;129-Gt(ROSA)26Sor^tm1(cre/ERT)Nat^/J (8 weeks old, 20-25g, male and female, R26CreER, strain # 4847 [RRID:IMSR_JAX:004847]) mice were purchased from The Jackson Laboratory (Bar Harbor, Maine, USA). Vegfr-1^flox^ and R26CreER were bred to generate tamoxifen-inducible Vegfr-1 knockout mice (Vegfr-1^flox/flox^R26CreER^+/-^, Vegfr-1^flox/flox^R26CreER^+/+^), and littermate controls (Vegfr-1^flox/flox^R26CreER^-/-^). All mice were housed in standard clear cages with free access to food and water. Controlled temperature (21°±1°C) and light/dark cycle of 12/12h. In all experiments, the investigators were blinded to groups and treatments, including data acquisition, sample processing, and data analyses. The animals were under isoflurane anesthesia (3% v/v in O_2_) for endometriosis induction, and tail snip sampling for genotyping. Carbon dioxide (CO_2_) inhalation was used for euthanasia. All efforts were made to minimize the number of animals and their suffering.

### Ethics statement

All experiments involving animals were conducted according to the ethical policies and procedures approved by the ethics committee of Boston Children’s Hospital Institutional Animal Care and Use Committee (IACUC, protocol number 19-12-4054R) and were in accordance with the International Association for Study of Pain (IASP) and ARRIVE 2.0 guidelines.

All experiments involving human samples were also approved by Institutional Review Boards (IRB). For samples from the Adolescence to Adulthood (A2A) cohort, the approval was conceived by the Boston Children’s Hospital IRB on behalf of Boston Children’s Hospital and Brigham and Women’s Hospital. Informed consent was obtained, with parental consent/participant assent for girls <18 years. The collection of additional samples was approved by the Mass General Brigham IRB under protocol 2017P000184. Importantly, in both cases, sampling was performed only after patients signed consent.

### Drugs and antibodies

Tamoxifen (Sigma-Aldrich, cat# T5648), Entrectinib (InvivoChem, Libertyville, IL, USA, CAS 1108743-60-7, cat# V0609); a mouse version of the rat anti-mouse VEGFR IgG1antibody MF1, (a kind gift from Eli Lilly & Co). anti-NGF (provided by Cygnal Therapeutics, Cambridge, MA), anti-BDNF (provided by Cygnal Therapeutics), IgG control (anti-L1CAM provided by Cygnal Therapeutics).

### Generation of VEGFR1 knockout mice

VEGFR1 knockout was induced by tamoxifen treatment in Vegfr-1^flox/flox^R26CreER^+/-^ and Vegfr-1^flox/flox^R26CreER^+/+^ mice. Littermate controls (Vegfr-1^flox/flox^R26CreER^-/-^) also received the treatment. Mice were treated via oral gavage with Tamoxifen (6 mg/mouse/150 μL of corn oil) every other day for 10 consecutive days. Endometriosis was induced 7 days after the last tamoxifen administration. VEGFR1 KO was confirmed by genotyping and immunohistochemical staining of the dorsal root ganglia (DRG).

### Induction of endometriosis

A non-surgical model of endometriosis-associated pain was induced as previously described^26^. The mice were acclimatized at least one week before the experiment began. Donor mice received a subcutaneous (s.c.) injection of 3 μg/mouse estradiol benzoate in sesame oil to stimulate endometrium growth. Four to seven days later, the uteri from donor mice were dissected and placed into a Petri dish containing Hank’s Balanced Salt Solution (HBSS, Thermo Fisher Scientific, Waltham, MA, USA). The uterine horns were split longitudinally with the aid of scissors. The horns from each donor mouse were minced in consistent fragments smaller than 1 millimeter (mm). Each recipient mouse received a dissociated uterine horn in 500 µL of HBSS intraperitoneally for endometriosis induction. One donor mouse was used for every two endometriosis mice. Sham mice received 500 µL of HBSS intraperitoneally.

### Experimental design

### Neurotrophic molecule signaling blockade

We used two approaches to block VEGFR1 signaling in endometriosis: i) anti-VEGFR monoclonal antibody and ii) VEGFR1 knockout (KO). Mice bearing endometriosis were treated with a monoclonal antibody anti-VEGFR or IgG isotope (mMF1, 45 mg/kg) subcutaneously (s.c.), twice a week starting 29 days post induction (29-56 d.p.i.). Mechanical hyperalgesia was assessed weekly. In the last week of treatments (49 – 56 d.p.i.) spontaneous abdominal behavior and thermal discomfort were quantified. On the 56^th^ day mice were euthanatized, and lesion size was determined. In another set of experiments, VEGFR1 conditional knockout was performed in CreER^+^ mice by tamoxifen administration (6 mg/animal) p.o. gavage every other day for 10 days. To account for the effects of VEGFR1 in the tissue of donor and recipient mice, endometriosis was induced in KO and littermate controls using uterine horns from KO or littermate control donor mice, totaling 4 experimental groups. After endometriosis induction, mechanical hyperalgesia was determined weekly using von Frey filaments. On the 56^th^ d.p.i., lesions and DRG were harvested for VEGFR1 KO confirmation by immunohistochemistry.

To target additional neurotrophin signaling two strategies were used: i) immunotherapy using neutralizing antibodies anti-NGF and anti-BDNF, and ii) pharmacological treatment with entrectinib a pan-selective inhibitor of tropomyosin receptor kinase (Trk) A, B, and C. Mice with endometriosis were treated with antibodies anti-NGF, anti-BDNF, or IgG isotope (10 mg/kg) subcutaneously (s.c.), twice a week beginning on day 29 post induction (29-56 d.p.i.). Mechanical hyperalgesia was assessed weekly. In the last week of treatments (49 – 56 d.p.i.) spontaneous abdominal behavior and thermal discomfort were quantified. Similarly, entrectinib was administered by oral gavage in different treatment schedules to total 60 mg/kg/week per group. Specifically, group 1 received entrectinib at 15 mg/kg every other day, group 2 at 20 mg/kg three times a week, and group 3 at 60 mg/kg once a week. On the 56^th^ day mice were euthanatized, and lesion size was determined. In the last week of treatments (49 – 56 d.p.i.) thermal discomfort was assessed for all groups, while spontaneous abdominal behavior was quantified in group 3 (T3) which received a single weekly treatment with 60 mg/kg. On the 56^th^ day, lesions, blood, and femur were harvested for lesion size determination, liver and kidney toxicity, and bone health assessment, respectively.

### Determination of VEGFR1 ligand levels

The concentrations of VEGFA, VEGFB, PlGF, and soluble VEGFR1 (sFL-1) were determined by Ella Automated Immunoassay System according to manufacturer’s instructions (ProteinSimple, Bio-Techne, Minneapolis, MN, USA). VEGFR1 occupancy was calculated according to ligand-receptor affinity, as previously described^29–31^. The measurement of VEGF from mouse endometriosis lesions was performed according to manufacturer’s instructions using mouse VEGF quantikine ELISA Kit (Cat# MMV00, R&D systems, Minneapolis, MN, USA).

### Immunostaining

For VEGFR1 immunohistochemistry, the mouse dorsal root ganglia (DRG) were dissected 56 days after endometriosis induction and post-fixed in 4% paraformaldehyde in phosphate buffered saline (PBS) (m/v) for 24h at room temperature (RT). Samples were dehydrated, paraffin embedded, and sectioned in a microtome (Harvard Medical School Rodent Histology Core). The 7 μm thick sections were deparaffinized and hydrated before antigen retrieval in citrate buffer was done. Slides were heated in a microwave until they reached 90°C and were cooled to RT. Endogenous peroxidase was inactivated with 3% hydrogen peroxide in methanol (v/v) for 15 min at RT. Sections were blocked in 3% bovine serum albumin (BSA) in PBS 0.5% triton-X 100 (m/v/v) for 1h at RT. Samples were incubated with rabbit anti-mouse VEGFR1 primary antibody (Abcam, Cambridge, UK, cat# 32152, 1:200 dilution in PBS-T [RRID:AB_778798]) overnight at 4°C. Slides were washed and incubated with goat anti-rabbit-HRP secondary antibody for 30 min at RT (Vector Laboratories, Newark, CA, USA, cat# MP-7451 [RRID:AB_2631198]). Color was developed using HRP substrate kit (Vector Laboratories, cat# SK-4105 [RRID:AB_2336520]) for 1 min and 45 seconds at RT. Slides were washed and counterstained with Gills III Formulation hematoxylin for 6 seconds, washed and dehydrated before slide mounting with Permount^TM^ mounting medium (Fisher Scientific, Waltham, MA, USA, cat# SP15-100).

For immunofluorescence analysis, mouse DRG and endometriotic lesions were dissected and post-fixed in 4% paraformaldehyde in phosphate buffered saline (PBS) (m/v) for 24h at 4°C. Samples were then dehydrated with 30% sucrose in PBS (m/v) for 48h at 4°C, followed by 30% sucrose in PBS and optimum temperature cutting reagent (OCT) (1:1, v/v) for 24h at 4°C. DRG and lesions were frozen in OCT, sectioned in a cryostat (16 μm thick), and placed on silanized slides. Sections were hydrated with PBS, blocked in 5% BSA in PBS 0.5% triton-X 100 (m/v/v) for 1h at RT, following overnight incubation at 4°C with primary antibodies: mouse anti-mouse phosphorylated-NF-κB (pNF-κB, Santa Cruz Biotechnology, Dallas, TX, USA, cat# sc-136548, 1:200 [RRID:AB_10610391]), rabbit anti-mouse TrkA (Invitrogen, Waltham MA, USA, cat. # MA5-32123, 1:100 [RRID:AB_2809414]), rabbit anti-mouse NGF (Abcam, Boston, MA, USA, cat. # AB52918, 1:300 [RRID:AB_881254]), and β-III tubulin (Novus Biologicals, Centennial, CO, USA, cat. # NB120-11314, 1:200 [RRID:AB_792496]). Slices were then incubated with appropriate secondary antibody: goat anti-mouse Alexa Fluor 594 secondary antibody (1:500, cat. # A21125, Thermo Fisher Scientific [RRID:AB_141593]), goat anti-rabbit Alexa Fluor 647 secondary (1:500, cat. # A-21235, Thermo Fisher Scientific [RRID:AB_2535804]), or goat anti-rabbit Alexa Fluor 488 (1:500, cat. # A-11008, Thermo Fisher Scientific [RRID:AB_143165]). DAPI was used to stain nuclei (Cayman Chemicals, Ann Arbor, MI, USA, cat. # 14285). Images were aquired and processed on a confocal microscope using 20x objective (Zeiss LSM 880 laser scanning microscope with Airyscan, Carl Zeiss Microscopy, Thornwood, NY, USA).

### Animal behavior

### Evoked abdominal mechanical hyperalgesia

Mechanical hyperalgesia was assessed as previously described^26^. Briefly, mice were habituated to the conditions for at least 2h for three consecutive days before pain assessment. Pain response to a mechanical stimulus (mechanical hyperalgesia) was measured using von Frey filaments. Abdominal mechanical hyperalgesia was determined by a trained experimenter. The area of external genitalia and consecutively stimulation in the same region were avoided. Jump or paw flinches were considered as a withdrawal response^32^. The up-and-down method was used to determine the mechanical threshold. Testing started with the 0.4g filament and measurements were calculated using a modified version of the open-source software Up-Down Reader^33^.

### Thermal gradient

The thermal gradient assay was performed as previously described^26^. Mice were placed on a metallic base catwalk with a continuous temperature gradient (7–50 °C). Animals walked freely while being video-recorded from above (Bioseb, France). Each run lasted 1.5 hours. After an exploration period (30 minutes), the time mice spent in each temperature zone was recorded. Data were presented as the time spent (in seconds) in each temperature zone during the last 60 minutes.

### Spontaneous pain behaviors

Spontaneous abdominal pain was quantified using abdominal licking, stretching (abdominal contortions), and squashing of the lower abdomen against the floor as previously described^26^. Direct abdominal licking was quantified for 10 minutes using bottom-up video recording as the number of times (bouts) the mouse directly groomed the abdominal region without going for any other region before or after the behavior. To measure abdominal contortions, mice were placed in individual chambers and the number of abdominal contortions was quantified over 10 minutes. The computed behaviour consisted of abdomila muscle contraction in combination with hind limbs streach. Abdominal squashing, was quantified during 5 minutes, and the considered behaviour was the number of times a mouse pressed its lower abdominal region against the floor. In all testing, the investigators were blinded to the groups and treatments.

### Liver and kidney toxicity determination

On the 56^th^ d.p.i., mice were euthanatized, and their blood was collected into heparinized collection tubes by cardiac puncture. Plasma was separated by centrifugation (3200 rpm, 10 min, 4°C). Kidney or liver toxicity was determined by the levels of urea, alanine aminotransferase (ALT), and aspartate aminotransferase (AST) in the plasma. The levels were determined by commercial kinetics kits according to manufacturer’s instructions at the Longwood Small Animal Imaging Facility (LSAIF) Blood lab at Beth Israel Deaconess Medical Center.

### Micro-computerized tomography analysis (μCT)

The right femur was collected 56 days after endometriosis induction from entrectinib or vehicle treated mice. Samples were fixed in paraformaldehyde 4% in PBS (m/v) for 24 h and maintained in 70% ethanol (v/v) until μCT analysis. The samples were scanned on a Bruker SkyScan 1173 microtomograph (Bruker BioSpin Corporation, Kontich, Belgium). NRecon, DataViewer, CTVox, and CTAn software was used to reconstruct and measure the images. The parameters analyzed after image acquisition were bone surface, volume, density, total porosity, and open-pore volume.

### Statistical analysis

Results are presented as mean ± SEM. The data were analyzed using GraphPad Prism version 8 (GraphPad Software, San Diego, CA, USA). Mechanical hyperalgesia was analyzed by two-way repeated measure analysis of variance (ANOVA), followed by Tukey’s *post hoc*. One-way ANOVA followed by Tukey’s *post hoc* was used to analyze data from experiments with a single time point. Comparison between two groups was conducted using Student’s t-test.

## Results

Classical neurotrophins signal by binding to tyrosine kinase receptors to induce neurite ingrown. The number of neurotrophic molecules and their receptor has steadily expanded since the discovery of NGF and TrkA and VEGF signaling via VEGFR1 has been shown to have similar effects. Therefore, we sought to evaluate the extent to which key neurotrophic molecules contribute to endometriosis-associated pain.

### Levels of VEGFR1 ligands are increased in the peritoneal fluid of endometriosis patients and in mouse lesions

The role of VEGF signaling through VEGFR2 to induce angiogenesis has long been recognized as a key means of supporting endometriosis lesion growth ^18,20,25,34,35^. However, the potential for VEGFR1 activation to support neurite ingrowth has only been appreciated relatively recently in the cancer context and has not been explored in endometriosis. Therefore, we first measured the levels of VEGFR1 agonists in the peritoneal fluid of patients undergoing endometriosis surgery **(Fig. 1 A-B)**. We found that the concentrations of VEGFA, VEGFB, PlGF, and sVEGFR1 were abundant in most evaluated samples **(Fig. 1B)**. Based on the levels of VEGFR1 agonists, the calculated VEGFR1 occupancy is sufficient to elicit neurite ingrowth and thereby pain **(Fig. 1C)**. Importantly, in peritoneal fluid, VEGFA concentrations were the major driver of VEGFR1 occupancy, with log[VEGFA] predicting 90% of the variance.

**Figure 1.**
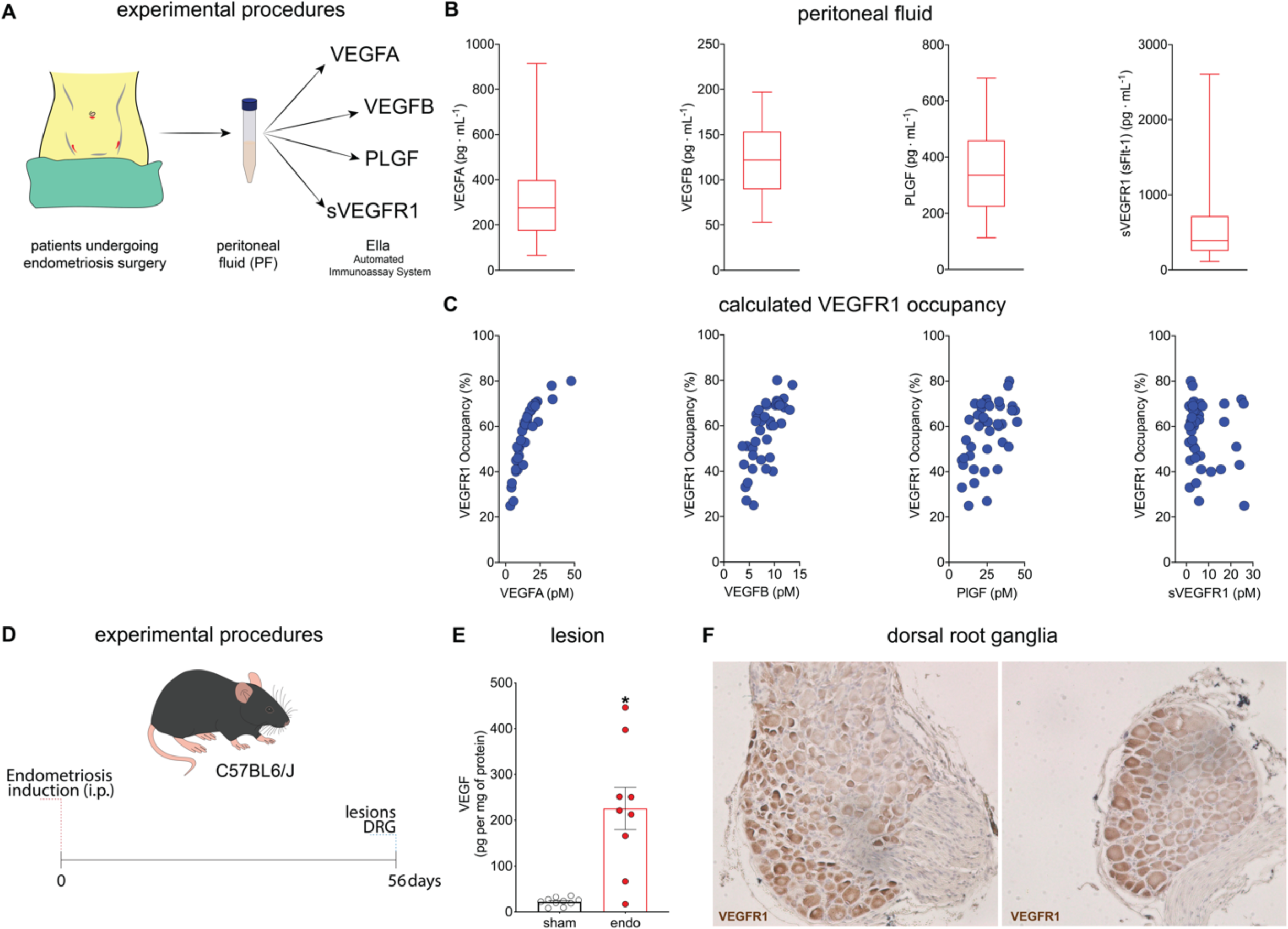
Levels of VEGFR1 ligands are increased in the peritoneal fluid of endometriosis patients and in mouse lesions. (A) experimental procedures for peritoneal fluid collection of endometriosis patients. (B) The levels of VEGFA, VEGFB, PlGF, and sVEGFR1 in peritoneal fluid samples determined by Ella^®^. Data are presented in box and whisker charts representing the levels of each mediator in pg/mL. (C) Calculated VEGFR1 occupancy. Note that in published data, occupancy is approximately proportional to signalling^65^. (D) Experimental procedures and timepoints for tissue collection in our mouse model. (E) 56 days after endometriosis induction, mouse lesions were collected for determination of VEGF levels by ELISA. Results are presented as mean ± SEM of VEGF levels, n = 10 (sham – uterine horn tissue), and n = 9 (endo) mice per group (*P < 0.05 vs. sham). (F) expression profile of VEGFR1 expression in DRG nociceptors determined by immunohistochemistry 56 days after endometriosis induction.

To determine whether VEGFR1 signaling might also be present in our mouse model, we next measured VEGFA levels in the mouse lesions. We demonstrated that VEGFA levels are increased in endometriotic lesions **(Fig. 1E)** and that its receptor VEGFR1 is consistently expressed in DRG neurons in naïve and endometriosis-bearing mice **(Fig. 1F)**. Therefore, based in previous published work ^16^ and the present evidence, we hypothesized that VEGFR1 signaling might be important in endometriosis-associated pain.

### VEGF neutralization or VEGFR1 ablation does not reduce endometriosis-associated pain or thermal discomfort in mice

To determine the extent to which VEGFR1 signaling is important for endometriosis-associated pain, we targeted this signaling using two different strategies. First, we measured the extent to which VEGFR1 neutralization with an anti-VEGFR1 monoclonal antibody (MF-1) would reduce endometriosis-associated pain in mice. We found that treatment with anti-VEGFR1 did not alter endometriosis-induced mechanical hyperalgesia **(Fig. 2B left)** or lesion size **(Fig. 2B right)**. Since spontaneous pain is the main complaint of patients with endometriosis, we next determined whether treatment with MF-1 would reduce endometriosis-associated spontaneous pain. We found that none of the spontaneous abdominal-related behaviors (licking, squashing, or contortions) were reduced by the treatment **(Fig. 2C)**. Finally, we assessed the mice’s own determination of discomfort using the thermal gradient assay. We confirmed that sham mice prefer temperatures 27 to 36 ^°^C with a stronger preference for 34 ^°^C, indicating that sham-treated mice were most comfortable at that temperature. In contrast, mice with endometriosis exhibited a more dispersed pattern ranging from 22 to 36 ^°^C with no single preferred temperature, suggesting that lesion-bearing mice were not comfortable at any temperature^26^. In corroboration with our previous results, blocking VEGF-VEGFR1 signaling did not reverse the loss of a comfort zone caused by lesions, as demonstrated by the dispersed occupancy pattern in MF-1-treated mice **(Fig. 2D).** Since we observed that treatment with anti-VEGFR1 did not reduce pain, we next wanted to establish the extent to which VEGFR1 is required for the incitement of pain. Therefore, we first developed a conditional tamoxifen-induced cre-dependent knockout mouse line **(Fig. 2E and F)**. VEGR1 floxed mice and littermate controls (LM) were treated with tamoxifen to induce cre-dependent VEGFR1 gene ablation **(Fig. 2F)**. Ablation of VEGFR1 expression was confirmed by immunohistochemistry in the dorsal root ganglia (DGR) **(Fig. 2G)**. We performed a four-group experiment using VEGFR1-floxed and LM control mice as donors or recipients to determine the extent to which the lack of VEGFR1 in donor tissue (mimicking retrograde menstruation) or in a combination of donor and recipient play a role in endometriosis lesion formation and pain **(Fig. 2H left)**. None of the investigated scenarios of VEGFR1 depletion demonstrated an analgesic effect **(Fig. 2H right)**. Therefore, while levels of VEGFR1 ligands are increased (both in humans and mice), our data indicate that neither VEGFR1 in donor nor from recipient mice contribute to endometriosis-associated pain.

**Figure 2.**
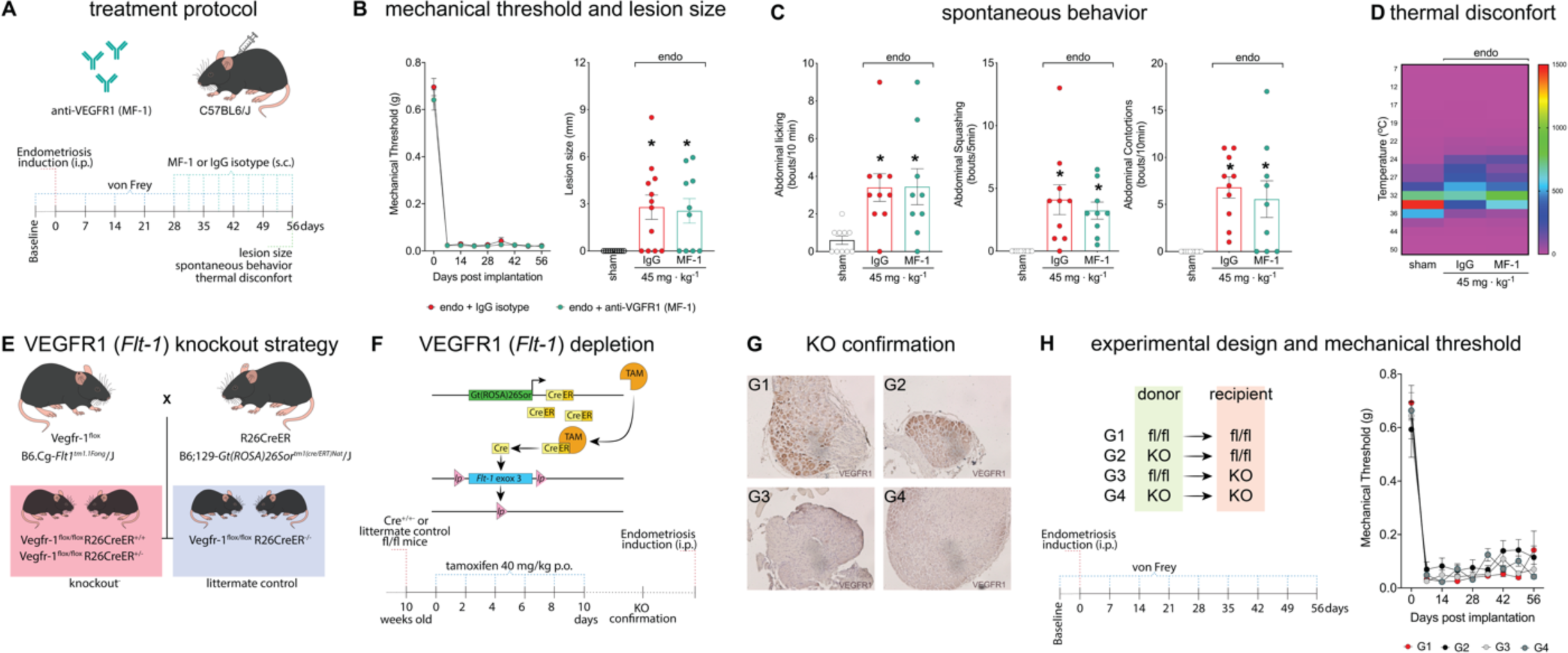
VEGF neutralization or VEGFR1 ablation do not reduce endometriosis-associated pain or thermal discomfort in mice. (A) scheme of the treatment protocol with anti-VEGFR1 (MF-1) antibody and IgG control. (B) mechanical hyperalgesia was determined using von Frey filaments before (zero) and after (7, 14, 21, 28, 35, 42, and 56 days) endometriosis induction. Results are presented as mean ± SEM of mechanical threshold, n = 10 mice per group. (C) spontaneous behaviours measurements. For abdominal licking, the total number of times that mice directly groomed the abdominal region (without going to any other body region before or after the behaviour) was quantified for 10 minutes. For abdominal squashing, the number of times the mice pressed the lower abdominal region against the floor was quantified for 5minutes. Sham mice did not display abdominal squashing. Abdominal contortions were quantified for 10 minutes by counting the number of contractions of the abdominal muscle together with stretching of hind limbs. Sham mice did not display abdominal contortions Results are expressed as mean ± SEM of abdominal licking, squashing, and contortions bouts per minute, n = 9-12. (*P < 0.05 vs. sham). (D) thermal discomfort heatmap. Heatmap shows mean time spent in each temperature zone for IgG control- or MF-1-treated mice. Data are presented as mean ± SEM of the amplitude of permanence in seconds in each thermal zone during 60 min, n = 9-12. (E) breeding scheme for generation of VEGFR1 tamoxifen-induced cre-dependent knockout strategy. (F) scheme of tamoxifen-induced cre-dependent VEGFR1 knockout and tamoxifen treatment protocol. (G) DRG neuron representative images for VEGFR1 knockout confirmation determined per IHC analysis. G1-G2 from littermate controls and G3-G4 from knockout mice n = 10. (H) Endometriosis induction protocol scheme with receptor and donor combinations. Mechanical response before (zero) and after (7, 14, 21, 28, 35, 42, and 56 days) endometriosis induction using von Frey filaments. Results are presented as mean ± SEM of mechanical threshold, n = 10 mice per group.

### Entrectinib, a pan-Trk inhibitor, reduces endometriosis-associated pain and thermal discomfort in mice

Patients with endometriosis have higher levels of NGF in lesion as well as genetic variations that correlates NGF to pain^25^ indicating that NGF might play an important role in human endometriosis pathology. Having found that VEGFR1 signaling does not contribute to pain in our model and because NGF is often found in endometriosis lesions, we next sought to investigate classical neurotrophins. To that end, we employed entrectinib, an inhibitor of the neurotrophin receptors TrkA, TrkB, and TrkC. Entrectinib reduced mechanical hyperalgesia **(Fig. 3B left)** and lesion size **(Fig. 3B right)** in all selected treatment schedules (T1 – T3) from day 42 to 56 post endometriosis induction. Weekly delivery of entrectinib at 60 mg/kg was proven more effective in restored comfort as measured in the thermal gradient when compared to vehicle treated mice **(Fig. 3C)**. Since reduced and more frequent doses showed a milder reduction to thermal discomfort (T1 and T2), the weekly dose of 60 mg/kg was chosen for the next experiments **(Fig. 3C)**. Because spontaneous pain is the main symptom among patients with chronic pain, we determined the effect of weekly treatment with entrectinib in spontaneous pain. We found that entrectinib-treated mice showed reduced endometriosis-associated spontaneous behaviors **(Fig. 3D)**. Altogether, our data show that disruption in neurotrophin-Trk signaling is effective in reducing endometriosis-associated pain.

**Figure 3.**
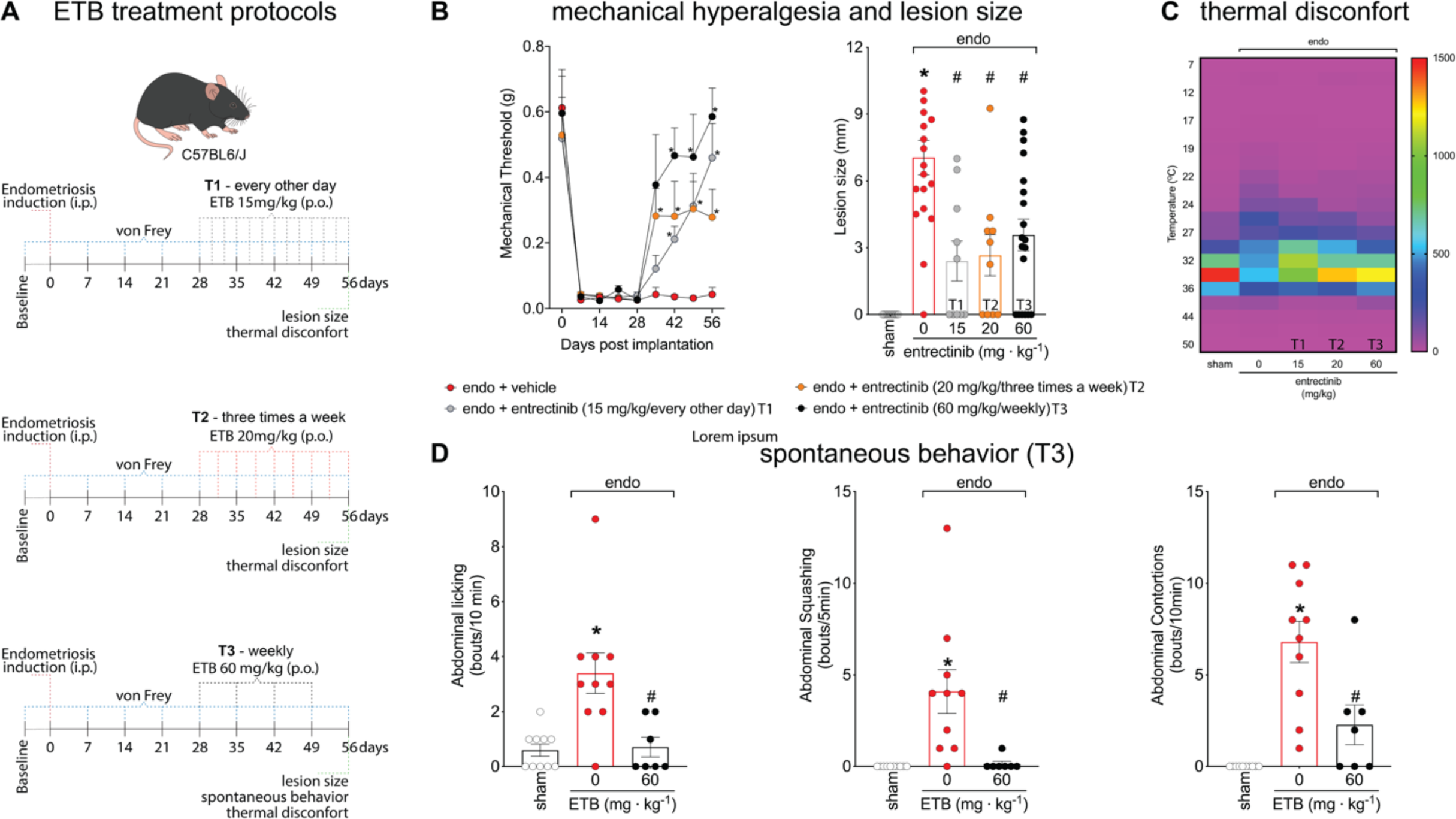
Entrectinib, a pan-Trk inhibitor, reduces endometriosis-associated pain and thermal discomfort. (A) scheme of the treatment protocol with entrectinib. T1: 15 mg/kg every other day; T2: 20 mg/kg three times a week; and T3: 60 mg/kg weekly. In all schedules of treatment, the maximum weekly dose is 60 mg/kg. (B left) mechanical response before (zero) and after (7, 14, 21, 28, 35, 42, and 56 days) endometriosis induction using von Frey filaments. Results are presented as mean ± SEM of mechanical threshold, n = 10 mice per group (*P < 0.05 vs. vehicle-treated group). (B right) lesion size. Results are presented as mean ± SEM of lesion size in mm, n = 10-19 mice per group (*P < 0.05 vs. sham, # P<0.05 vs. vehicle-treated group). (C) thermal discomfort heatmap. Data are presented as mean ± SEM of the amplitude of permanence in seconds in each thermal zone during 60 min. (D) spontaneous behaviour measurements. For abdominal licking, the total number of times that mice directly groomed the abdominal region (without going to any other body region before or after the behaviour) was quantified for 10 minutes. For abdominal squashing, the number of times the mice pressed the lower abdominal region against the floor was quantified for 5 minutes. Sham mice did not display abdominal squashing. Abdominal contortions were quantified for 10 minutes by counting the number of contractions of the abdominal muscle together with stretching of hind limbs. Sham mice did not display abdominal contortions. Results are expressed as mean ± SEM of abdominal licking, squashing, and contortions bouts per minute, n = 7-10. (*P < 0.05 vs. sham, # P < 0.05 vs. vehicle-treated group).

### Neutralizing NGF, but not BDNF, reduces endometriosis-associated pain and thermal discomfort in mice

To determine the contribution of the Trk ligands NGF (TrkA) and BDNF (TrkB) in our validated mouse model of endometriosis-associated pain, we then disrupted neurotrophin receptor signaling. For that, we used neutralizing antibodies against NGF and BDNF. Upon measuring mechanical hyperalgesia, lesion size, spontaneous behaviors, and thermal discomfort **(Fig. 4A)**, we observed that anti-NGF immunotherapy substantially reduced mechanical hyperalgesia from the 42^nd^ to 56^th^ days after endometriosis induction **(Fig. 4B left)**; other treatments were indeffective. Although analgesic effects were maintained up to 56 d.p.i. in mice treated with anti-NGF, no differences were observed in lesion size **(Fig. 4B right)**. Based on these results, we next analyzed spontaneous pain-associated behaviors and thermal discomfort only using anti-NGF therapy **(Fig. 4C-D)**. We found that the treatment with anti-NGF reduced endometriosis-induced abdominal licking **(Fig. 4C left)** and squashing **(Fig. 4C middle)**. A decrease in abdominal contortions was also observed, however the difference vs. IgG-treated mice did not reach statistical significance **(Fig. 4C right)**. In the thermal gradient assay **(Fig. 4D)**, anti-NGF therapy reduced thermal discomfort, restoring the amplitude of time spent in specific thermal zones, when compared to isotype-treated control mice **(fig. 4D)**. Altogether, our results show that NGF-TrkA signaling, but not BDNF-TrkB contributes to endometriosis-associate pain.

**Figure 4.**
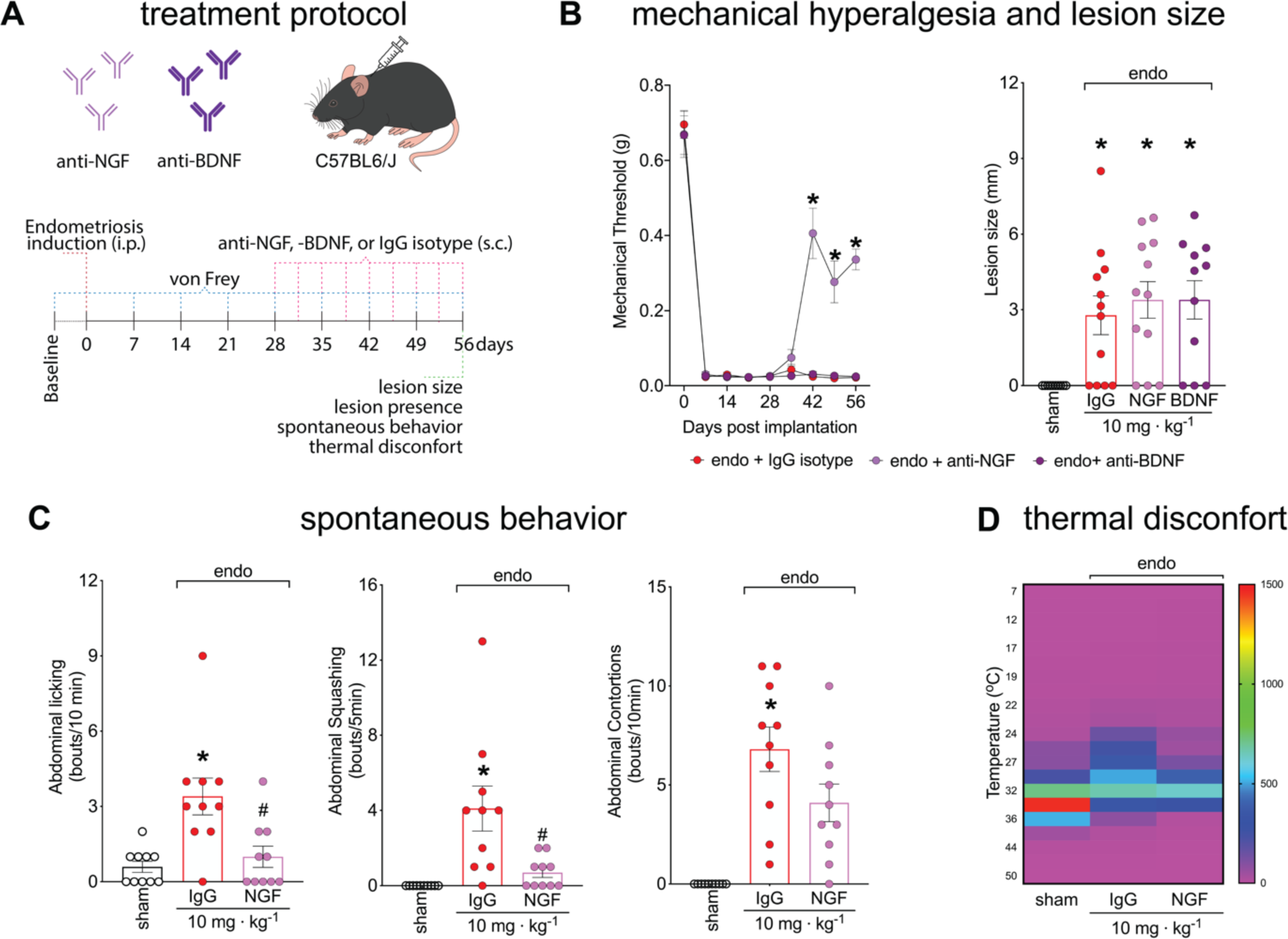
NGF, but not BDNF neutralization, reduces endometriosis-associated pain and thermal discomfort in mice. (A) scheme of the treatment protocol with IgG control, anti-NGF, or anti-BDNF antibodies. (B) mechanical hyperalgesia was measured using von Frey filaments before (zero) and after (7, 14, 21, 28, 35, 42, and 56 days) endometriosis induction. Results are presented as mean ± SEM of mechanical threshold, n = 10-12 mice per group (*P < 0.05 vs. IgG treated group). (C) spontaneous behaviours measurements. For abdominal licking, the total number of times that mice directly groomed the abdominal region (without going to any other body region before or after the behaviour) was quantified for 10 minutes. For abdominal squashing, the number of times the mice pressed the lower abdominal region against the floor was quantified for 5minutes. Sham mice did not display abdominal squashing. Abdominal contortions were quantified for 10 minutes by counting the number of contractions of the abdominal muscle together with stretching of hind limbs. Sham mice did not display abdominal contortions. Results are expressed as mean ± SEM of abdominal licking, squashing, and contortions bouts per minute, n = 10 mice per group. (*P < 0.05 vs. sham, # P < 0.05 vs. IgG control). (D) thermal discomfort heatmap. Heatmap shows mean time spent in each temperature zone for IgG control-, anti-NGF-, or anti-BDNF-treated mice. Data are presented as mean ± SEM of the amplitude of permanence in seconds in each thermal zone during 60 min, n = 10 mice per group.

### NGF-TrkA signaling is activated during endometriosis

To confirm that NGF’s role in endometriosis pathogenesis^25^ is accurately reflected in our mouse model **(Fig. 5A)**, we used immunofluorescence to stain for NGF in mouse lesions. Interestingly, nociceptors seem to be closely located to this NGF gradient, as observed by colocalization between NGF and β-tubulin III (TUJ3), a pan-neuronal marker **(Fig. 5B)**. In the DRG, lesion bearing mice displayed a higher percentage of TrkA-expressing neurons, and most importantly, we found that TrkA^+^ nociceptors from mice with endometriosis demonstrated increased activation as observed higher percentage of TrkA^+^pNF-κB^+^ in comparison to sham **(Fig. 5C bottom, D)**. Altogether, these data suggests that NGF-TrkA signaling is activated and might contribute to pain in our model of endometriosis.

**Figure 5.**
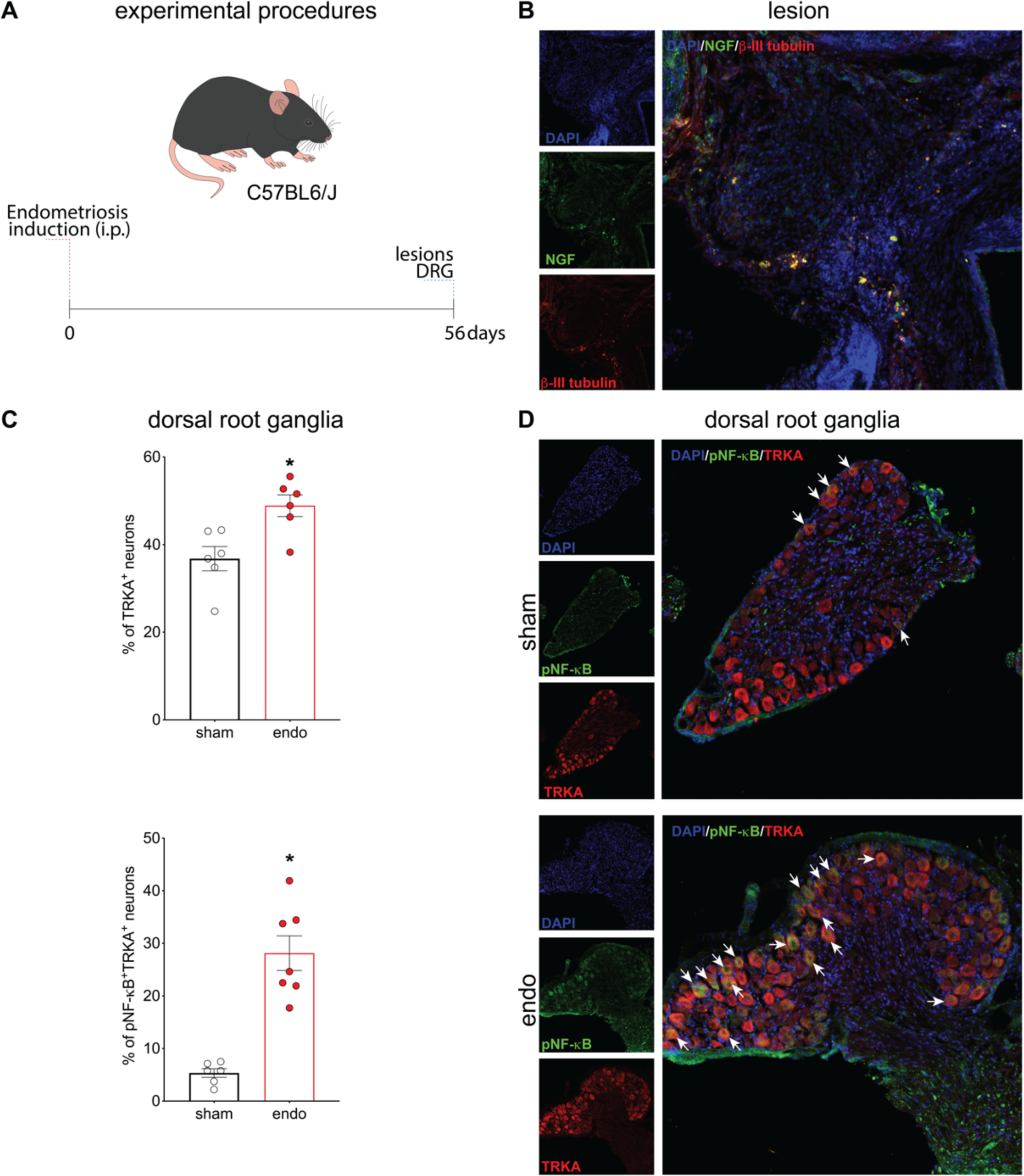
NGF-TrkA signaling is activated during endometriosis. (A) scheme of experimental procedures. (B) representative image from endometriotic lesions stained for NGF and beta-III tubulin. (C) quantification of TrkA^+^ and pNF-κB^+^TrkA^+^ neurons in dorsal root ganglia (DRG) of sham and endometriosis lesion-bearing mice. Lesions and DRG were dissected at 56 dpi. Results are presented as mean ± SEM of the percentage of positive neurons. n = 6 or 7 mice per group. (*P < 0.05 vs. sham). (D) Representative images of DRG neurons stained for TrkA (red) and p-NF-kB (green) by confocal microscopy.

### Weekly treatment with entrectinib does not induce weight change, liver or kidney toxicity, or bone loss in mice

As mentioned, clinical trials with entrectinib in children and young adults demonstrated a high incidence of bone fractures (clinicaltrials.gov, NCT02650401)^36^. Given the generalized analgesic effect of entrectinib with the different treatment schedules **(Fig. 3A, T1, T2, and T3)**, we next wanted to determine its safety. Even though the weekly dose proposed here is much lower than the ones used in clinical trials, even after human equivalent dose determination, we sought to determine kidney and liver function, and bone loss in mice **(Fig. 6A)**. Weekly treatment with entrectinib did not induce weight changes **(Fig. 6B)**, nor kidney or liver function alteration as per levels of urea **(Fig. 6C)**, alanine aminotransferase (ALT) **(Fig. 6D Left)**, or aspartate transaminase (AST) **(Fig. 6D right)** in the plasma, respectively. Moreover, weekly treatment did not alter bone parameters in the femur, as determined per micro-computerized tomography analysis **(Fig. 6E)**. No significant changes were observed in femur surface, volume, density, or porosity in the evaluated dose and schedules of treatment. We found, however, that increasing number of treatments, such as every other day **(Fig 6, T1)** or three times a week **(Fig 6, T2),** reduced bone porosity **(Fig. 6E)**. This indicates that while increased number of treatments with lower doses have analgesic effects, this substantially affects bone porosity. This indicates that increased treatment schedules, rather than dose, might be a limiting factor for entrectinib use.

**Figure 6.**
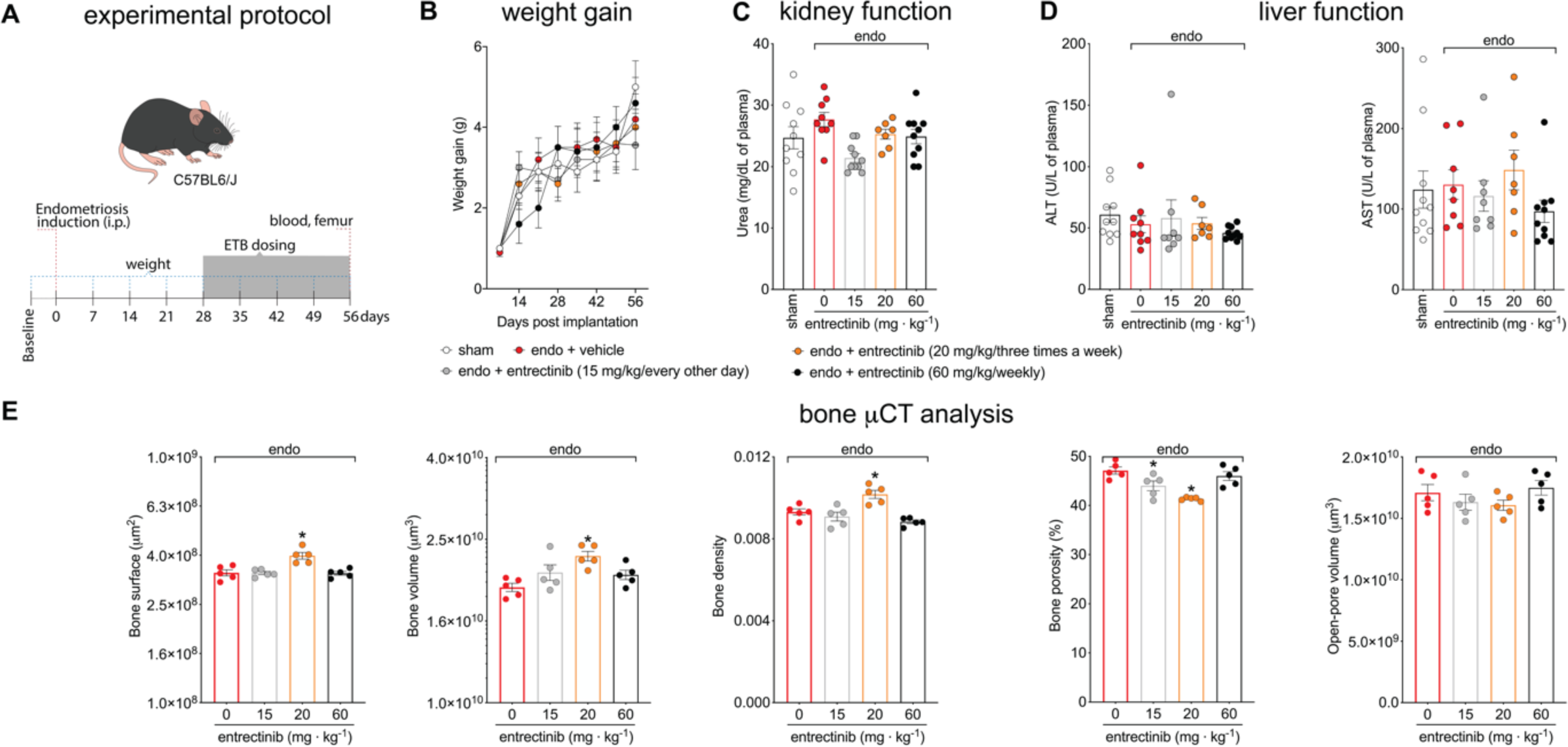
Weekly treatment with entrectinib does not induce weight change, liver or kidney toxicity, or bone loss in mice. (A) scheme of experimental protocol for determination of entrectinib safety using the different treatment schedules. (B) mouse weight was determined weekly. Results are presented as mean ± SEM in grams, n = 7-10 mice per group. (C) kidney function was determined by measuring urea plasma levels. Results are presented as mean ± SEM of urea levels, n = 7-10 mice per group. (D) liver function was determined by measuring ALT and AST plasma levels. Blood and femur were collected 56 dpi (after 4 weeks of treatment). Results are presented as mean ± SEM of AST and ALT levels, n = 7-10 mice per group. (E) Microcomputed tomography of bone (femur) was used to determine bone loss. Results are presented as mean ± SEM, n = 5 mice per group.

## Discussion

Pain is one of the key presenting symptoms of endometriosis. While pain correlates poorly with lesion characteristics or rASRM-defined disease stage^37^, peri-lesional TRPV1 staining correlates with chronic pelvic pain^38,39^ and women with endometriosis-associated pain exhibit dramatically increased nerve fiber densities^40,41^. Lesion microvessel density is correlated with pain^42^, suggesting that factors controlling both may play an important role in neurite recruitment. NGF is a key neurotrophic factor^25^ to maintaining TRPV1 neuronal expression^43,44^ and has long been known to induce angiogenesis *in vivo* ^45^. In addition, when this work began, VEGF-A, acting through VEGFR1 had recently been shown to modulate pain in the context of cancer by supporting neuronal recruitment^46^.

Prior to our work, it had already been shown that NGF is present at higher concentrations in the peritoneal fluid of endometriosis patients vs. controls and that this results in increased neurotrophic activity^47^. It was also well-established that VEGF-A is upregulated in lesion tissue^48–54^ and peritoneal fluid^55–64^, but the expression of other VEGFR1 ligands had not been explored. We observed VEGFA, VEGFB, PlGF, and sVEGFR1 in the peritoneal fluid of patients undergoing endometriosis surgery at levels that are more than sufficient to induce VEGFR1 activation and signaling^65^ and that VEGFA predominated in occupancy calculations. In our validated mouse model^26^, we found high levels of VEGF in the lesions as well as the presence of its receptor in primary nociceptor neurons, suggesting a possible signaling axis to pain sensitivity. However, blocking VEGFR1 signaling with anti-VEGFR1 antibody or cKO did not reduce pain or lesion size. This indicates that while present in the lesion and peritoneal fluid, VEGF-VEGFR1 signaling does not mediate pain in our model. Importantly, this model results in spontaneous pain, evoked pain, and discomfort and these are alleviated by drugs known to alleviate pain in human patients^26^. These data suggest that anti-VEGFR1 treatment is unlikely to be effective in treating endometriosis.

In contrast, we demonstrated that NGF/TrkA, but not BDNF/TrkB signaling, contributes to endometriosis-related spontaneous and evoked pain responses. Blocking NGF signaling with entrectinib, a pan-Trk inhibitor, reduced evoked abdominal mechanical pain as well as spontaneous pain and discomfort. Specifically, we demonstrated that endometriosis-associated pain is mediated via NGF-TrkA signaling, since blocking NGF with an antibody reduced pain to approximately the same extent as entrectinib, while blocking BDNF signaling had no effect on pain, notwithstanding the expected effect on mouse body weight. The fact that anti-NGF and entrectinib treatment had similar effect sizes also suggests, but does not prove, that TrkC ligands may have little effect in our model.

This observation is consistent with literature showing that neurotrophic factors, such as NGF, participate in inflammatory pain^66^ and neuropathic pain^67^. NGF contributes both indirectly and directly to nociceptor neuron sensitization and pain. NGF signaling via TrkA in immune cells (e.g., mast cells, basophils, macrophages)^68–70^ results in the release of NGF and other pro-nociceptive molecules, such as interleukin (IL)-1β^71^. NGF also directly induces nociceptor neurons sensitization and activation^15^. Specifically for endometriosis, a recent GWAS study highlighted that variance among genes such as NGF is associated with pain presentation^25^. Therefore, we hypothesized that neurotrophins (*e.g.,* NGF and BDNF) participate in endometriosis-associated pain. Our data show that NGF is co-localized with TUJ3 in endometriosis and that TrkA^+^ DRG neurons are activated during endometriosis. This corroborates human findings that show both NGF and TrkA are highly expressed in endometriosis lesions and are correlated with nerve fiber density and deep dyspareunia^47^. In corroboration, we found that anti-NGF immunotherapy reduced endometriosis-associated evoked and spontaneous pain behaviors in mice. Anti-NGF treatment reduced mechanical hyperalgesia, abdominal licking, squashing, and contortions, and decreased thermal discomfort. Similarly, in a model of cyclophosphamide-induced cystitis, treatment with anti-NGF reduces peripheral hypersensitivity in mice^72^. Pain inhibition was also observed in the same model of cystitis in rats when animals were treated with the NGF sequestering protein REN1820^73^. Humanized anti-NGF monoclonal antibodies have been clinically tested in osteoarthritis^74–77^, low back pain^78^, diabetes-associated neuropathy^79^, and interstitial cystitis^80^, corroborating the observed phenomena. On the other hand, we found that anti-BDNF immunotherapy did not reduce any of the evaluated parameters. Altogether, this indicates that NGF-TrkA signaling, but not BDNF-TrkB, mediates endometriosis-associated pain in our model. Herein, we demonstrated weekly treatments with low doses of entrectinib (60 mg/kg) reduced evoked and non-evoked pain behaviors in mice. This suggests that discontinuous disruption of NGF-TrkA signaling, by weekly treatments with entrectinib, is sufficient to decrease endometriosis-associated pain. This effect is likely to be via NGF-TrkA signaling since pain (evoked and spontaneous) in our model was reduced after treatment with anti-NGF, but not anti-BDNF. In this context, it is worth noting that NGF targeting therapies are often linked to undesired side effects. Of greatest concern, while all studies show that anti-NGF decreases pain, in a portion of patients with osteoarthritis, anti-NGF therapy was correlated to joint destruction and the need for total joint replacement^25^. Similarly, treatments with entrectinib in children and young adults increased the incidence of bone fractures (clinicaltrials.gov, NCT02650401)^36^. Based on the clinical relevance of NGF-targeting treatment for endometriosis pain management, we addressed the safety of entrectinib in the dose and schedule of treatments tested in this study. Herein, we demonstrated that weekly treatment with entrectinib at 60 mg/kg did not induce changes in weight gain, liver, or kidney toxicity, nor bone morphology. Therefore, in this pre-clinical study, we found that discontinuous low doses of entrectinib reduced endometriosis-associated pain while not showing significant side effects.

## Conclusion

In this study we demonstrated that while VEGFR1 agonists are upregulated in the peritoneal fluid of women with endometriosis, blocking this signaling does not reduce pain in a murine model. On the other hand, we found that the neurotrophin NGF (but not BDNF) is key for endometriosis-associated pain. Blocking this signaling with anti-NGF or entrectinib is effective at reducing abdominal mechanical pain, spontaneous pain, and thermal discomfort in our model. Moreover, weekly treatments with entrectinib did not show significant side effects in mice. The lack of drug efficacy at reducing ongoing pain drives most endometriosis therapy failure. Therefore, our study pinpoints NGF as a key factor in endometriosis-associated pain and establishes NGF-TrkA signaling as a potential target for the development of novel therapies for endometriosis.

## Funding

This work was supported by grants from The J. Willard and Alice S. Marriott Foundation and Marriott Daughters Foundation.

## Acknowledgments

T.H.Z. acknowledges international split PhD scholarship (*Programa de Doutorado Sanduíche no Exterior* – PDSE) from Coordination for the Improvement of Higher Education Personnel (CAPES, Brazil, finance code 001). We also thank the support of Multiuser Center for Research Laboratories of Londrina State University for the access to µCT equipment free of charge. W.A.V.J. acknowledges CNPq fellowship (#309633/2021-4). We are also thankful for the collaboration of Rachel Arredondo for English editing this manuscript.

## Declaration of interests

The authors declare no conflicts of interest.

## Notes

### Competing Interest Statement

The authors have declared no competing interest.

